# Modulation of brain activity by selective attention to audiovisual dialogues

**DOI:** 10.1101/781344

**Authors:** Alina Leminen, Maxime Verwoert, Mona Moisala, Viljami Salmela, Patrik Wikman, Kimmo Alho

## Abstract

In real-life noisy situations, we can selectively attend to conversations in the presence of irrelevant voices, but neurocognitive mechanisms in such natural listening situations remaiin largely unexplored. Previous research has shown distributed activity in the mid superior temporal gyrus (STG) and sulcus (STS) while listening to speech and human voices, in the posterior STS and fusiform gyrus when combining auditory, visual and linguistic information, as well as in left-hemisphere temporal and frontal cortical areas during comprehension. In the present functional magnetic resonance imaging (fMRI) study, we investigated how selective attention modulates neural responses to naturalistic audiovisual dialogues. Our healthy adult participants (N = 15) selectively attended to video-taped dialogues between a man and woman in the presence of irrelevant continuous speech in the background. We modulated the auditory quality of dialogues with noise vocoding and their visual quality by masking speech-related facial movements. Both increased auditory quality and increased visual quality were associated with bilateral activity enhancements in the STG/STS. In addition, decreased audiovisual stimulus quality elicited enhanced fronto-parietal activity, presumably reflecting increased attentional demands. Finally, attention to the dialogues, in relation to a control task where a fixation cross was attended and the dialogue ignored, yielded enhanced activity in the left planum polare, angular gyrus, the right temporal pole, as well as in the orbitofrontal/ventromedial prefrontal cortex and posterior cingulate gyrus. Our findings suggest that naturalistic conversations effectively engage participants and reveal brain networks related to social perception in addition to speech and semantic processing networks.

## 1 Introduction

In everyday life, we are often faced with multiple speaker situations, for instance, when dining in a crowded restaurant or talking to a friend while hearing a radio in the background. Such situations require segregation of speech streams originating from different sources and selection of one of the streams for further processing. The neural mechanisms through which this type of attentional selection is achieved are not yet fully understood (e.g., Rimmele et al., 2015).

A meta-analysis (Alho et al., 2014) of functional magnetic resonance imaging (fMRI) studies on stimulus-dependent sound processing and attentional related modulations in the auditory cortex showed that speech and voice processing activate overlapping areas in the mid superior temporal gyrus and sulcus bilaterally (STG and STS, respectively). Furthermore, selective attention to continuous speech appeared to modulate activity predominantly in the same areas (Alho et al., 2014). Importantly, selectively attending to a particular speaker in a multi-talker situation results in the STG activity that represents the spectral and temporal features of attended speech, as if participants were listening only to that speech stream (Mesgarani and Chang, 2012). In other words, the human auditory system restores the representation of an attended speaker while suppressing irrelevant or competing speech.

In addition to STG/STS, selective attention to non-speech sounds engages prefrontal and parietal cortical areas (Alho et al., 1999, Degerman et al., 2006, Tzourio et al., 1997, Zatorre et al., 1999), which has been associated with top-down control needed to select attended sounds and reject irrelevant sounds. Selective attention to continuous speech, however, does not appear to markedly engage prefrontal and superior parietal areas (Alho et al., 2003, Alho et al., 2006, Scott et al., 2004). This is likely because selective listening to speech is a highly automatized process, less dependent on fronto-parietal attentional control (Alho et al., 2006; see also Mesgarani and Chang, 2012). Such automaticity might be due to listeners’ lifelong experience in listening to speech. However, initial orienting of attention to one of three concurrent speech streams has yielded enhanced activation in the fronto-parietal network, hence, purportedly engaging an attentional top-down control mechanism (Alho et al., 2015, Hill and Miller, 2010).

Natural situations with multiple speakers might not only be complicated by a demand to listen selectively to one speech stream while ignoring competing speech, but also by degraded quality of the attended speech (e.g., when talking in a noisy café on the phone with a poor signal). Studies addressing the comprehension of degraded (e.g., noise-vocoded) speech involving only one speech stream have reported increased activity in the posterior parietal cortex (Obleser et al., 2007) and frontal operculum (Davis and Johnsrude, 2003) as compared to more intelligible speech. Listening to degraded, yet intelligible and highly predictable speech, in turn, elicits activity in the dorsolateral prefrontal cortex, posterior cingulate cortex, and angular gyrus (e.g., Obleser et al., 2007). Moreover, the amount of spectral detail in speech signal was found to correlate with STS and left inferior frontal gyrus (IFG) activity, regardless of semantic predictability (Obleser et al., 2007). McGettigan and colleagues (2012) observed increasing activity along the length of left dorsolateral temporal cortex, in the right dorsolateral prefrontal cortex and bilateral IFG, but decreasing activation in the middle cingulate, middle frontal, inferior occipital, and parietal cortices associated with increasing auditory quality. Listening to degraded speech has also activated the left IFG, attributed to higher-order linguistic comprehension (Davis and Johnsrude, 2003) and the dorsal fronto-parietal network, related to top-down control of attention (Obleser et al., 2007). Overall, increased speech intelligibility enhances activity in the STS (McGettigan et al., 2012, Obleser et al., 2007, Scott et al., 2000), STG (Davis and Johnsrude, 2003), middle temporal gyrus (MTG; Davis and Johnsrude, 2003), and left IFG (Davis and Johnsrude, 2003, McGettigan et al., 2012, Obleser et al., 2007). Enhanced activity in these areas is therefore may be related to enhanced speech comprehension with increasing availability of linguistic information.

The studies described above, however, used only single-speaker paradigms. Recently, Evans and colleagues (2016) examined how different masking sounds are processed in the human brain. They used a selective attention paradigm with two speech streams, namely, a masked stream and a target stream. The target speech was always clear, whilst the masked speech was either clear, spectrally rotated or noise-modulated. Increased intelligibility of the masked speech activated the left posterior STG/STS, however, less extensively than a clear single speech alone. This was taken to suggest that syntactic and other higher order properties of masking speech are not actively processed and the masker sounds may be actively suppressed already at early processing stages (see also Mesgarani and Chang, 2012). In contrast, the masked speech yielded increased activation in the frontal (bilateral middle frontal gyrus, left superior orbital gyrus, right IFG), parietal (left inferior and superior parietal lobule) and middle/anterior cingulate cortices, as well as in the frontal operculum and insula. These activations were suggested to reflect increased attentional and control processes. The results corroborate those from earlier positron emission tomography (PET) studies (e.g., Scott et al., 2004) on selective attention to a target speaker in the presence of another speaker (speech-in-speech) or noise (speech-in-noise). More specifically, Scott and colleagues (2004) found more activity in the bilateral STG for speech-in-speech than speech-in-noise, whereas speech-in-noise elicited more activity in the left prefrontal and right parietal cortex than speech-in-speech. Scott and colleagues suggested that these additional areas might be engaged to facilitate speech comprehension or that they are related to top-down attentional control. Correspondingly, Wild and colleagues (2012) reported activations in frontal areas (including the left IFG) that were only present when the participants selectively attended to the target speech among non-speech distractors. In contrast to studies reporting increased left IFG activations to increased intelligibility of degraded speech (Davis and Johnsrude, 2003, McGettigan et al., 2012, Obleser et al., 2007), Wild and colleagues (2012) found greater activity in the left IFG for degraded than for clear target speech. By contrast, STS activity was increased with decreasing speech intelligibility, regardless of attention. Increased activity for attended degraded speech was proposed to reflect “the improvement in intelligibility afforded by explicit, effortful processing, or by additional cognitive processes (such as perceptual learning) that are engaged under directed attention” (Wild et al., 2012, p. 14019). The authors further suggested that top-down influences on early auditory processing might facilitate speech comprehension in difficult listening situations.

The majority of fMRI studies on selective attention to speech have used only auditory speech stimuli (e.g., Alho et al., 2003, Alho et al., 2006, Evans et al., 2016, Puschmann et al., 2017, Wild et al., 2012). However, natural conversations often include also visual speech information. Integrating a voice with mouth movements (i.e., visual speech) facilitates speech understanding in relation to mere listening (Sumby and Pollack, 1954). In accordance, fMRI studies on listening to speech have shown that the presence of visual speech enhances activity in the auditory cortex and higher order speech-processing areas (e.g., Bishop and Miller, 2009, McGettigan et al., 2012). A related magnetoencephalography (MEG) study showed that the presence of visual speech enhances auditory-cortex activity that follows the temporal amplitude envelope of attended speech (Zion Golumbic et al., 2013; for similar electroencephalography (EEG) evidence, see O’Sullivan et al., 2015). Facilitation of speech comprehension by visual speech holds especially true for noisy situations (e.g., Sumby and Pollack, 1954) and degraded quality of attended speech (e.g., McGettigan et al., 2012, Zion Golumbc et al., 2013). Some fMRI studies have suggested maximal facilitation of speech comprehension by visual speech at intermediate signal-to-noise ratios of auditory information (McGettigan et al., 2012, Ross et al., 2007).

Degraded speech increases demands for fronto-parietal top-down control (Davis and Johnsrude, 2003, Evans et al., 2016), whereas adding visual speech appears to facilitate selective attention (Sumby and Pollack, 1954, Zion Golumbic et al., 2013). However, it is still unknown whether fronto-parietal areas are activated during selective attention to visually degraded speech. Moreover, an earlier study that employed a factorial design with different levels of auditory and visual clarity in sentences (McGettigan et al., 2012) did not include an unmodulated (clear) visual and auditory condition. Hence, to our knowledge, brain responses to continuous naturalistic dialogues with varying audio-visual speech quality have not been systematically examined before.

In the current study, we collected whole-head fMRI data in order to identify brain regions critical for selective attention to natural audiovisual speech. More specifically, we examined attention-related modulations in the auditory cortex and associated fronto-parietal activity during selective attention to audiovisual dialogues. In addition, we assessed an interplay between auditory and visual quality manipulations. We also included clear auditory and visual stimulus conditions to investigate brain areas activated during selective attention to naturalistic dialogues in the presence of irrelevant clear speech in the background. Our experimental setup might be regarded as mimicking watching a talk show on a TV while a radio program is playing on the background. Comparing brain activity during attention to the dialogues with activity during control conditions where the dialogues are ignored and fixation cross is to be attended, allowed us to determine attention-related top-down effects and distinguish them from stimulus-dependent bottom-up effects (Alho et al., 2014).

We predicted that both increased speech intelligibility and increased amount of visual speech information in the attended speech would be associated with stronger stimulus-dependent activity in the STG/STS as well as subsequent activity in brain areas involved in linguistic processing. Moreover, we hypothesized that degrading auditory or visual quality of attended speech might be related to increased fronto-parietal activity due to enhanced attentional demands. Finally, we were interested to see whether attention to audiovisual speech and the quality of this speech would have interactions in some brain areas involved in auditory, visual or linguistic processing, or in the control of attention.

## 2. Methods

### 2.2. Participants

Fifteen healthy right-handed adult volunteers (5 males, age range 20–38 years, mean 25.3 years) participated in the present study. All participants were native Finnish speakers with normal hearing, normal or corrected-to-normal vision, and no history of psychiatric or neurological illnesses. Handedness was verified by the Edinburgh Handedness Inventory (Oldfield, 1971). An informed written consent was obtained from each participant before the experiment. The experimental protocol was approved by the Ethics Review Board in the Humanities and Social and Behavioural Sciences, University of Helsinki.

### 2.2. Stimuli

### 2.1. Stimulus preparation

The stimuli consisted of 36 video clips showing scripted spoken dialogues (see Table 1 for an example of a dialogue). The topics of dialogues were of neutral, such as weather, vacation, and study plans. The grammatical structure of dialogues was matched as closely as possible, preserving their naturalness. An independent native Finnish speaker subsequently verified the neutrality of dialogues as well as their meaningfulness and grammaticality. Each dialogue always consisted of seven lines spoken alternatingly by two actors, and each line contained 9–13 words.

**Table 1.** Example of one natural speech dialogue by two actors (A and B) used in the experiment.

| Dialogue lines | Approximate English translation |
| --- | --- |
| A: Pitäisi kohta käydä kaupassa hakemassa välipalaa. Ostanko sinullekin jotain? | A: I should go soon to the store and get something to eat. Should I get something for you as well? |
| B: Ei kiitos tarvitse, minä pakkasin leipää ja jogurttia tänä aamuna lounaaksi. | B: No thanks, I packed bread and yoghurt with me for lunch today. |
| A: Hyvä on, mutta haluatko tulla mukaan seuraksi kauppaan kuitenkin? | A: Okay, but would you still like to come along with me to the store? |
| B: Mielelläni, voisin katsoa, jos löytäisin sieltä jotain syötävää myöhemmälle. | B: With pleasure, I could see if I would find something to eat later. |
| A: Haluatko tulla kanssani puistoon syömään, kun olemme tulleet kaupasta? | A: Would you like to come with me to the park to eat after visiting the store? |
| B: Ulkona on aika kylmä tänään. Mentäisiinkö mieluummin jonnekin sisälle? | B: It's quite cold outside today. Should we rather go somewhere inside? |
| A: Totta, voisimme siinä tapauksessa syödä täällä yliopiston kahvihuoneessa. | A: True, in that case we could eat here at the university in the coffee room. |

The stimulus recordings took place in a soundproof studio. The video clips were recorded with a wide angle (23.5 mm G lens) HXR-NX70E digital video camera (SONY Corporation, Tokyo, Japan). Two external microphones were attached to the camera in order to record the left and right audio channels separately (48 kHz sampling frequency, 16-bit quantization).

The actors were two native Finnish speakers (a male and female university student recruited for the recording purposes). They were unaware of the experimental setup and were compensated for their work. The actors memorized the dialogues beforehand but uttered their lines with a natural pace. An external prompter (programmed with Matlab version R2016, Mathworks Inc., Natick, MA, USA) was used to remind each actor to hold a pause before uttering the next line. The pause duration information was used in the subsequent fMRI data processing. The mean duration of dialogues was 60 s (range 55–65 s) with mean line duration of 5.4 s and inter-line pause duration of 3.4 s. Half of the dialogues started with the female speaker and the other half with the male speaker. The speakers sat next to one another with their faces slightly tilted towards each other, making the visual speech setting as natural as possible while maintaining visual speech information visible to a viewer. The video data were then edited with Corel VideoStudio Pro X 8 software (Corel Corporation, Ottawa, Ontario, Canada) and, finally, with Matlab (Mathworks Inc., Natick, MA, USA), see below. The video clips were cut into separate dialogues with 720 ms (18 frames) before the first and after the last spoken words. Thereafter, the videos were split into separate video and audio channels for subsequent editing with Adobe Audition CS6 (Adobe Systems Inc., San Jose, CA, USA) software. The audio channels were then converted to mono, cleaned from all non-voice background sounds, low-pass filtered at 5000 Hz, and scaled to have the same peak sound energy in all dialogues.

In addition to the natural speech, the audio data were noise-vocoded (Shannon et al., 1995, Davis and Johnsrude, 2003) using Praat software (version 6.0.27; Boersma and Weenink, 2001). The audio files were divided into 2 and 4 logarithmically spaced frequency bands between 300 and 5000 Hz (2 band cut-off points: 300, 1385, 5000 and 4 band cut-off points 300, 684, 1385, 2665, 5000). The filter bandwidths were set to represent equal distances along the basilar membrane (according to the Greenwood (1990) equation relating filter position to best frequency). The amplitude envelope from each frequency band was extracted using the standard Praat algorithm. The extracted envelope was then applied to band-pass filtered noise in the same frequency bands. Then, the resulting bands of modulated noise were recombined to produce the distorted speech. Noise vocoded speech sounds like a harsh robotic whisper (Davis and Johnsrude, 2003). Finally, the unchanged F0 (frequencies 0–300 Hz) was added to the noise-vocoded speech in order to maintain the speakers’ gender identity clearly perceivable and their voices distinguishable from the irrelevant voice speaking in the background (see below). The speech was perceived to be hardly intelligible with 3 frequency bands (i.e., 2 noise-vocoded bands and the intact F0 band) and quite intelligible with 5 frequency bands (i.e., 4 noise-vocoded bands and the intact F0 band). These two frequencyband manipulations for noise-vocoding were assumed to be optimal for our study on the basis of a behavioral pilot experiment. In this pilot experiment, 5 listeners (not included in the actual fMRI experiment) rated the intelligibility of seven dialogues noise-vocoded across a wide range of frequency bands (2, 4, 6, 8, 10, 12 or 16) with a non-vocoded F0 band. The participants listened to the dialogues one line at a time and provided a typed report on what they could hear. On average, for 2 and 4 noise-vocoded bands, 26.2 % (SD = 18.6) and 76.4 % (SD = 10.35) of the lines were perceived correctly.

In addition to manipulating auditory information, we parametrically varied the amount of visual speech seen by the participants. This was done by adding different amounts of dynamic white noise onto the region in the videos showing the speakers’ faces in Matlab R2016, using custom-made scripts. Five experienced viewers (not included in the actual fMRI experiment) confirmed that adding the noise reduced the visual quality so that the mouth movements and facial features were only poorly visible at highest noise level.

In the final step of stimulus preparation procedure, we recombined the ‘poor’, ‘medium’, and ‘good’ auditory quality sound files (with 2 noise-vocoded bands and an intact F0 band, 4 noise-vocoded bands and an intact F0 band, and clear intact speech, respectively) with the ‘poor’, ‘medium’, and ‘good’ visual quality video files (more masked poorly perceivable visual speech, less masked quite perceivable visual speech, and unmasked clear visual speech, respectively) video files using a custom-made Matlab script. The resulted videos were then compressed using VirtualDub software (http://www.virtualdub.org).

Taken together, each dialogue had 3 visual and 3 auditory quality variants, which resulted in altogether 9 experimental conditions, one for each quality combination (e.g., poor visual and good auditory quality) with three dialogues in each. All combinations were presented to the participants but each participant saw a different variant of each dialogue.

Furthermore, to increase the attentional load, we added continuous background speech as an auditory distractor. For this purpose, we chose a cultural history audio book (the Finnish translation of *The Autumn of the Middle Ages* by Johan Huizinga), which is freely distributed online by the Finnish Broadcasting Company (Yleisradio, YLE, https://areena.yle.fi/1-3529001). The book was read by a female professional Finnish-native actor. In order for the F0 in this auditory distractor to be approximately equidistant from the F0s of our female (200.4) and male (122.1) actors, we manipulated the F0 of the reader’s voice by using square root of the mean of the female and male voices in the recorded video clips. After some further manipulations based on the estimation of three experienced listeners, the resulting F0 was 156 Hz. The F0 manipulation was performed in Audacity software (https://sourceforge.net/projects/audacity/). The background speech was otherwise presented in its natural form and low-pass filtered at 5000 Hz to match the audio used in the experimental conditions. The audiobook was always presented as clear (i.e., non-vocoded) speech in the background. In addition, loudness differences between attended and unattended speech was kept minimal, as verified by three experienced listeners.

### 2.3. Procedure

Stimulus presentation was controlled through a script written in Presentation 20.0 software (Neurobehavioral Systems Inc., Berkeley, CA, USA). The video clips were projected onto a mirror mounted on the head coil and presented in the middle of the screen. All auditory stimuli were presented binaurally through insert earphones (Sensimetrics model S14; Sensimetrics, Malden, MA, USA). The experiment consisted of 3 functional runs with all 9 experimental conditions (Auditory Quality either poor, medium or good and Visual Quality either poor, medium or good) presented in each run along with 2 visual control conditions. The order of conditions was also randomized, however, the visual control conditions were always presented at the 6^th^ and 7^th^ place within a run. There was a small break of 40 s between these two dialogues. During the rest period, the participants were asked to focus on the fixation cross. Within all three functional runs, the order of the conditions was randomized for each participant. The competing audio distractor (audiobook) was presented 500–2000 ms before video onset and stopped at the offset of the video. The differing durations of dialogues were compensated for by inserting periods with a fixation cross between the instruction and the onset of the dialogue, keeping the overall trial durations constant.

#### 2.3.1. Attention-to-speech conditions

In attention-to-speech conditions, the participants were asked to attentively watch the videos, ignore the background speech, and after each 7-line dialogue answer to seven questions, one question related to each line of the dialogue. More specifically, they were instructed to answer whether a certain topic was discussed in a particular line (see Table 2) by pressing the “Yes” or “No” button on a response pad with their right index or middle finger, respectively. Each written question was presented on the screen for 2 s, during which the participant gave his/her answer. Regardless of the duration of the participant’s answer, the next question always started 2 s after the previous one. After the 7 questions, the participants were provided with immediate feedback on their performance (number of the correct answers), and the fixed duration of feedback was 2 s. Thereafter a next dialogue was presented after a short (2-s) written instruction on whether the task was an attention-to-speech or a control (see Section 2.3.2.) task and the fixation cross period (presented for 3–13 s to make all the trials equally long). All video clips had a rotating white fixation cross (inside a light grey box), placed in the middle of the screen, with a minimum of 9 and maximum of 15 rotations at a random interval but with minimum of 3 s between rotations. In the attention-to-speech condition, the participants’ task was to ignore the cross and concentrate on viewing the people speaking.

**Table 2.**
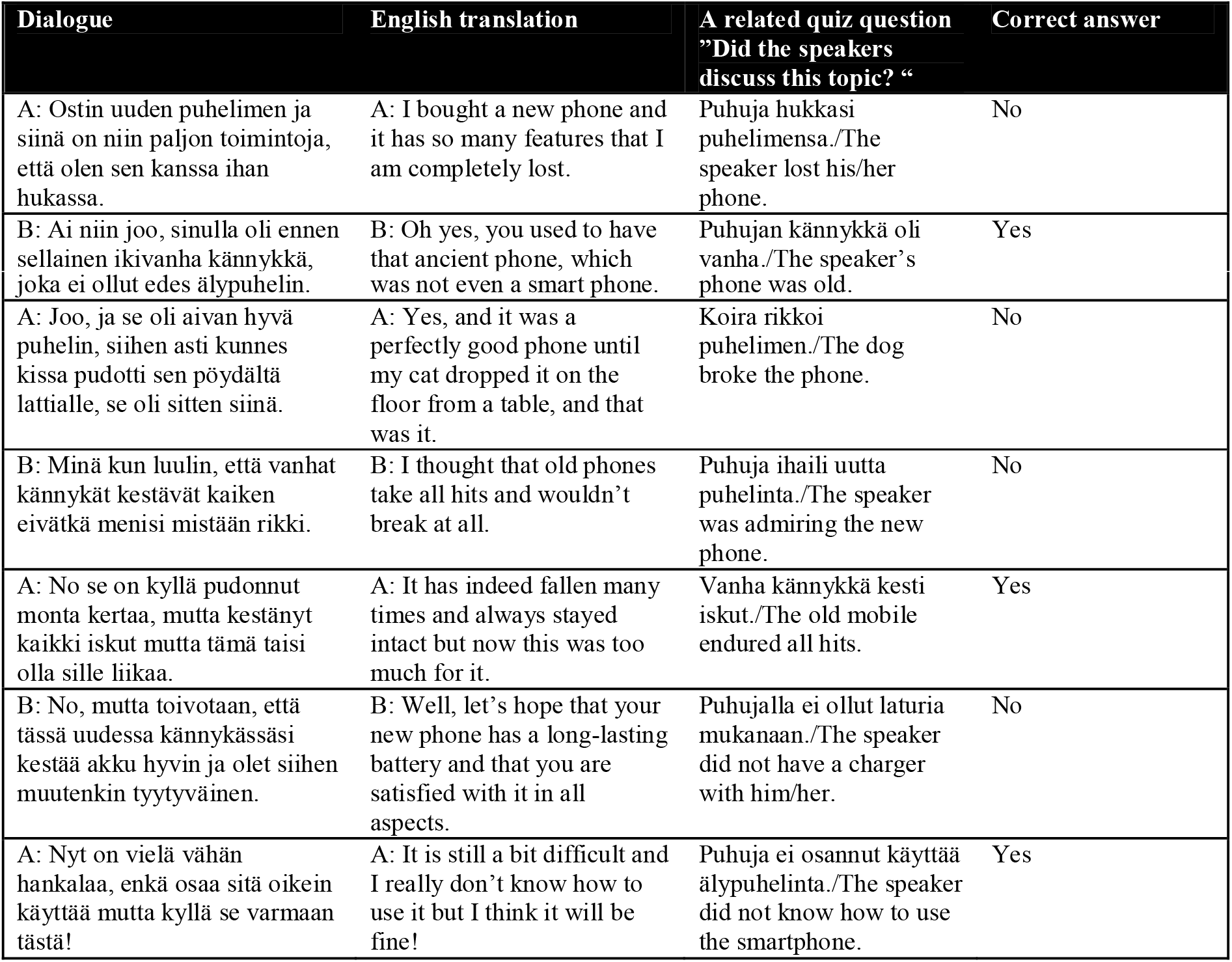
Example of quiz questions of a practice dialogue.

#### 2.3.2. Attention-to-the-fixation-cross condition

In addition to the attention-to-speech conditions, we included two control conditions. These consisted of videos with a combination of good auditory and good visual quality and a combination of poor auditory and poor visual quality. The dialogues in these control conditions were the ones not used in the attention-to-speech conditions. Identically to the attention-to-speech conditions, all video clips had a rotating white fixation cross (inside a light grey box), placed in the middle of the screen, with a minimum of 9 and maximum of 15 rotations from “×” to “+”, or vice versa, at random intervals but with a minimum of 3 s between the rotations. Thus, the attention-to-the-fixation-cross conditions used the same setup as the attention-to-speech conditions, but the task of the participants was to concentrate on counting the number of times the fixation cross rotated and ignore the dialogue and the background voice. After each control block, the participants were presented with seven questions (“Did the cross turn X times?” the X being 9, 10, 11, 12, 13, 14 or 15 in an ascending order), and they were asked to answer each question with “Yes” or “No” by pressing the corresponding button on the response pad with their right index or middle finger, respectively. Thereafter, the participants were provided with immediate feedback (e.g., “6/7 correct”).

#### 2.3.3. Practice trial

Before the actual fMRI scanning, the participants were familiarized with the task outside the scanner by viewing one practice dialogue with all conditions and answering questions related to its content.

### 2.4. Data acquisition

Functional brain imaging was carried out with 3T MAGNETOM Skyra whole-body scanner (Siemens Healthcare, Erlangen, Germany) using a 30-channel head coil. The functional echo planar (EPI) images were acquired with an imaging area consisting of 43 contiguous oblique axial slices (TR 2530 ms, TE 32 ms, flip angle 75°, voxel matrix 64 × 64, field of view 20 cm, slice thickness 3.0 mm, in-plane resolution 3.1 mm × 3.1 mm × 3.0 mm). Three functional runs of 368 volumes were measured for each participant. A total of 1158 functional volumes were obtained in one session (session duration approximately 50 min). High-resolution anatomical images (voxel matrix 256 × 256, in-plane resolution 1 mm × 1 mm × 1 mm) were acquired from each participant prior to the functional runs.

### 2.5. Data analysis

The fMRI data were pre-processed and analyzed in Statistical Parametric Mapping (SPM12; Wellcome Trust Centre for Neuroimaging, London, UK). The first 4 volumes in each run were dummies and were discarded in further analysis of the data, leaving 382 total volumes per run to be analyzed. The data were slice-time corrected, motion corrected, realigned to the middle image from each run, high-pass filtered (cutoff 1/260 Hz) and spatially smoothed with a Gaussian kernel of 6 mm. The images were normalized to MNI space using a standard pre-processing function in Conn software (Whitfield-Gabrieli & Nieto-Castanon, 2012). For the first-level statistical analysis, the general linear model was created including a regressor for each condition. Separate regressors were also included for (1) the instructions and the responses from the participant and (2) the quiz. This resulted in 13 regressors in total. Additionally, six movement parameters (3 translations, 3 rotations) were included as nuisance regressors. The conditions were modelled using a standard boxcar model. For the second-level analysis, we used the Multivariate and Repeated Measures (MRM) toolbox (McFarquhar et al., 2016). The contrast images of the nine experimental conditions compared to rest from each participant were entered into a 3 × 3 full factorial repeated-measures analysis of variance (ANOVA) with factors Visual Quality (3 levels: poor, medium, good) and Auditory Quality (3 levels: poor, medium, good). Within this model, F-contrasts were computed for the main effects and the interaction effect. A separate 2 × 2 repeated-measures ANOVA was conducted to account for stimulus quality and attentional effects. This additional ANOVA included factors Audiovisual Quality (2 levels: poor auditory and poor visual quality vs. good auditory and good visual quality) and Attention (2 levels: attention to speech vs. attention to the fixation cross). All reported contrasts were thresholded voxel-wise at *p* <. 001 with a cluster extend threshold of 100 voxels, resulting activity maps that were family-wise error (FWE) corrected at the cluster level, p(FWE) < 0.05. Statistical analyses of the responses to the quiz questions during the attention-to-speech condition were submitted to the repeated-measures ANOVA with factors Visual Quality (poor, medium, good) and Auditory Quality (poor, medium, good). For all analyses, the Greenhouse-Geisser correction was applied when the assumption of sphericity was violated. IBM SPSS Statistics 24 (IBM SPSS, Armonk, NY, USA) was used for conducting this analysis.

## 3. Results

### 3.1. Behavioral results

The mean performance scores for the attention-to-speech conditions are shown in Figure 1. Behavioral results demonstrated a significant main effect of Auditory Quality [F(2, 28) = 57.57, p = .001, η_p_^2^ = .80] and a significant main effect of Visual Quality [F(2, 28) = 8.2, p = .002, η_p_^2^ = .37]. Although visual quality appeared to have a slightly stronger effect on performance when the auditory quality was poor than when it was medium or good (see Figure 3) interaction between the two factors did not reach significance [F(4, 56) = 1.64, p = .176]. Bonferroni-corrected post-hoc tests for Auditory Quality revealed significant differences between all Auditory Quality conditions (for all comparisons, p < .001). Post-hoc tests for Visual Quality revealed significant differences between the poor and good quality conditions (p < .001) and between medium and good quality conditions (p = .029). The mean performance scores for the attention-to-the-fixation-cross conditions were 6.7/7 (SEM 0.16/7) and 6.5/7 (SEM 0.36/7) for the poor audiovisual quality condition and good audiovisual quality condition, respectively.

**Figure 1.**
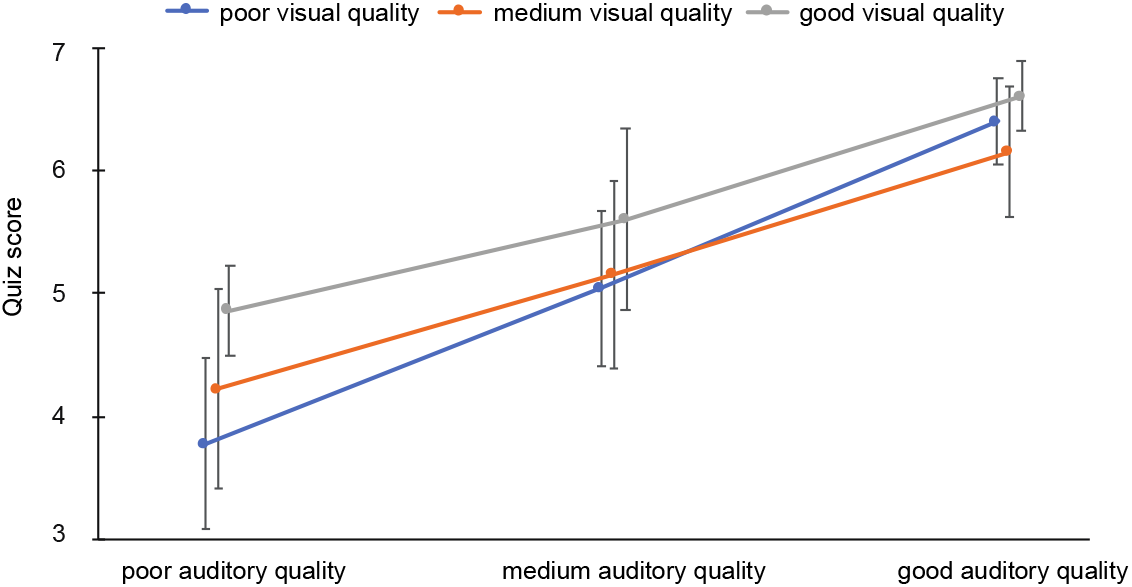
Mean performance scores (± SEM) as a function of Auditory Quality and Visual Quality. Chance level was 3.5.

### 3.2. fMRI results

#### 3.2.1 Auditory and visual quality

Figure 2 depicts the brain areas where the 3 × 3 ANOVA showed significant main effects of Auditory Quality (3 levels) and Figure 3 significant main effects of Visual Quality (3 levels) on brain activity measured during attention to speech. Figures 4 and 5 also depict mean parameter estimates of significant clusters, displaying the direction of the observed cluster effect. No significant interactions between these factors were found with the applied significance threshold.

**Figure 2.**
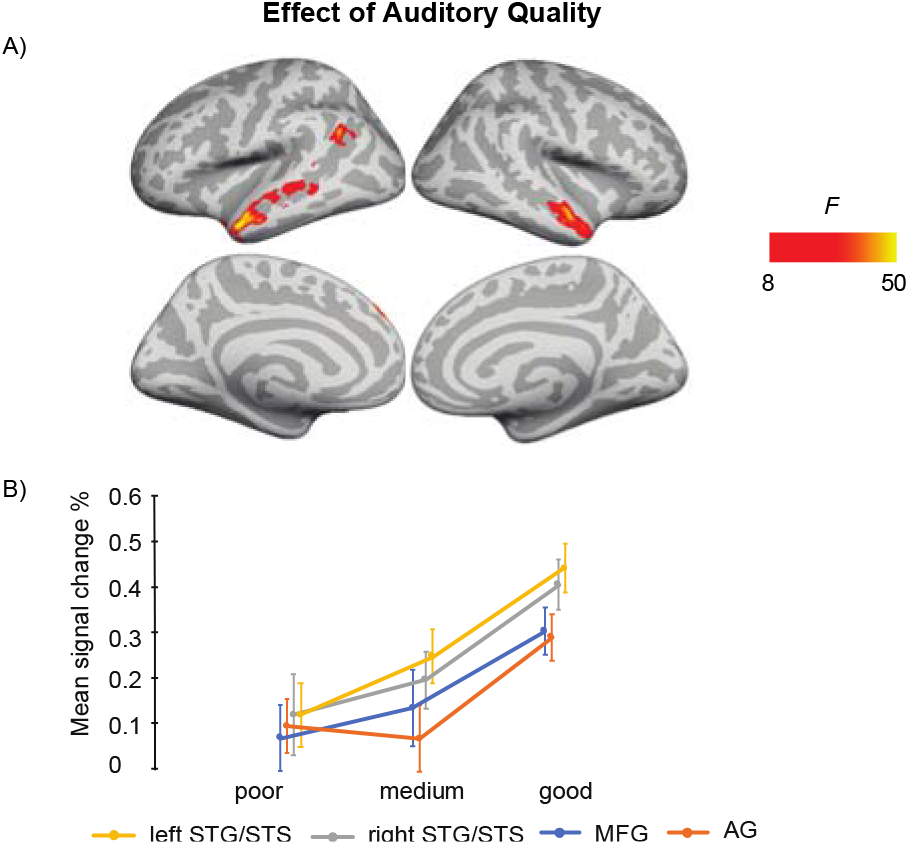
A: Brain areas showing significant main effects of Auditory Quality of attended audiovisual speech; p(cluster-level FWE) < .05, voxel-wise threshold p < .001, cluster extent k = 100. B: Mean signal changes (%) compared with rest in the clusters showing significant effects of Auditory Quality. STG, superior temporal gyrus; STS, superior temporal sulcus; AG, angular gyrus; MFG, middle frontal gyrus.

As seen in Figure 2, Auditory Quality showed a significant effect on brain activity in the STG/STS bilaterally, these effects extending from mid-STG/STS areas to the temporal poles, in the left angular gyrus, and the left medial frontal gyrus. Figure 4 demonstrates that in all these areas activity enhanced with increasing auditory quality. Note that due the projection of the clusters onto the cortical surface, the clusters in the superior temporal cortex are not seen as continuous in Figure 2.

Visual Quality, in turn, had significant effects on brain activity in the temporal and occipital cortices (Figure 3). As seen in Figure 3b, increasing visual quality was associated with enhancing activity in the STS bilaterally, this activity extending in the right hemisphere even to the temporal pole, and in the left inferior frontal gyrus. However, Figure 3c shows that in the left middle occipital cortex and the right fusiform gyrus, the activity was higher for poorer visual quality.

**Figure 3.**
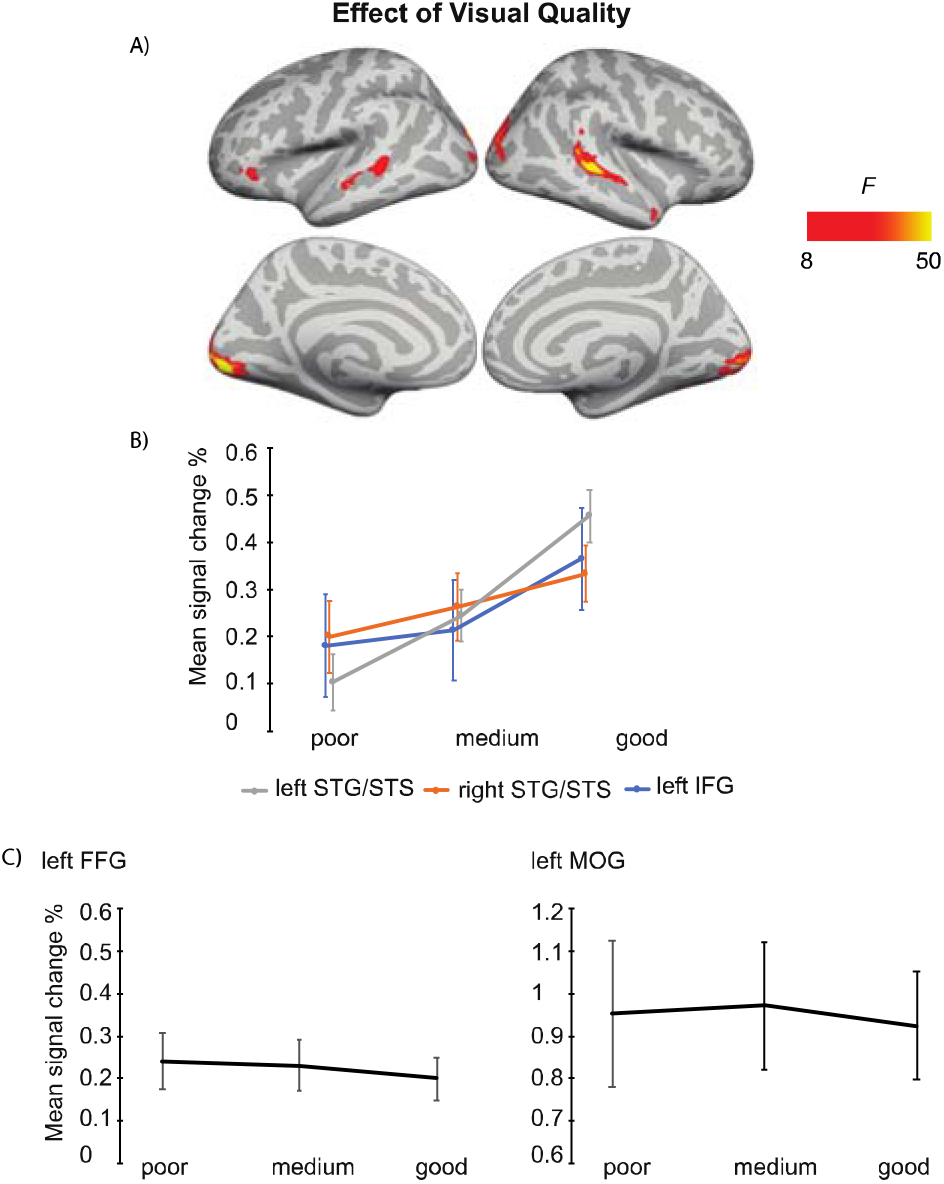
A: Brain areas showing significant main effects of Visual Quality of attended audiovisual speech; p(cluster-level FWE) < .05, voxel-wise threshold p < .001, cluster extent k = 100. B: Mean signal changes (%) compared with rest in the clusters showing significant effects of Visual Quality conditions and clusters showing increasing activity with increasing visual quality. C: The same for clusters showing decreasing activity with increasing visual quality. STG, superior temporal gyrus; STS, superior temporal sulcus; IFG inferior frontal gyrus; FFG, fusiform gyrus; MOG, middle occipital gyrus.

#### 3.2.2 Attention to speech versus attention to the fixation cross

Figure 4 depicts the brain areas where the 2 × 2 ANOVA showed significant main effects of Audiovisual Quality (2 levels: poor auditory and poor visual quality vs. good auditory and good visual quality). Figure 5 shows the brain areas where the 2 × 2 ANOVA showed significant main effects of Attention (2 levels: attention to speech vs. attention to the fixation cross) on brain activity. Figures 4 and 5 also depict mean parameter estimates of significant clusters, displaying the direction of the observed cluster effect. No significant interactions between the factors Attention and Audiovisual Quality were observed with the applied significance threshold. As seen in Figure 4, in the 2 × 2 ANOVA, there was a significant effect of Audiovisual Quality bilaterally in the STG/STS, these activations extending to the temporal poles, as well as in the left superior parietal lobule, left precuneus, the dorsal part of the right inferior parietal lobule, and in the middle occipital gyrus bilaterally. As seen in Figure 4b, in the left and right superior temporal gyri, the activity was enhanced with increasing audiovisual quality both during attention to the speech and attention to the fixation cross. In contrast, in both attention conditions, activity was higher in the left superior parietal lobule, the right inferior parietal, the left precuneus, and bilateral middle occipital gyrus for poorer audiovisual quality (Figure 4c).

**Figure 4.**
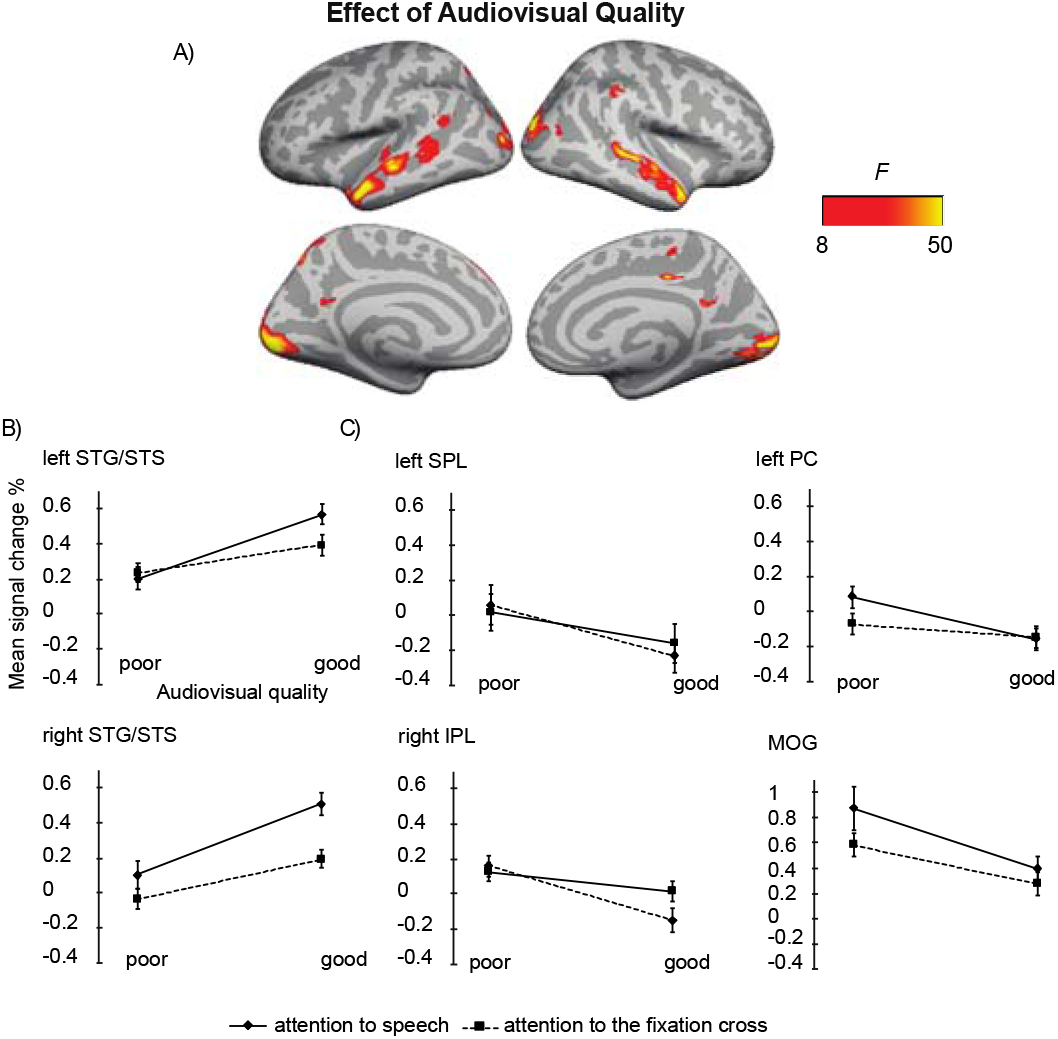
A: Brain areas showing significant main effects of Audiovisual Quality (good auditory and good visual quality vs. poor auditory and poor visual quality) across conditions with attention to speech and attention to the fixation cross; p(cluster-level FWE) < .05, voxel-wise threshold p < .001, cluster extent k = 100. B: Mean signal changes (%) compared with rest in the clusters showing significant effect of Audiovisual Quality and higher activity for better audiovisual quality. C. The same for clusters showing lower activity for higher audiovisual quality. STG, superior temporal gyrus; STS, superior temporal sulcus; IPL, inferior parietal lobule; SPL, superior parietal lobule; PC, precuneus; MOG, middle occipital gyrus.

Figure 5 demonstrates that Attention had a significant effect on brain activity in the left planum polare, the left angular gyrus, the left lingual gyrus, the right temporal pole, the right supramarginal gyrus, the right inferior parietal lobule, as well as in the ventromedial prefrontal cortex/orbitofrontal cortex and posterior cingulate bilaterally. In all these areas, activity was higher during attention to speech than during attention to the fixation cross.

**Figure 5.**
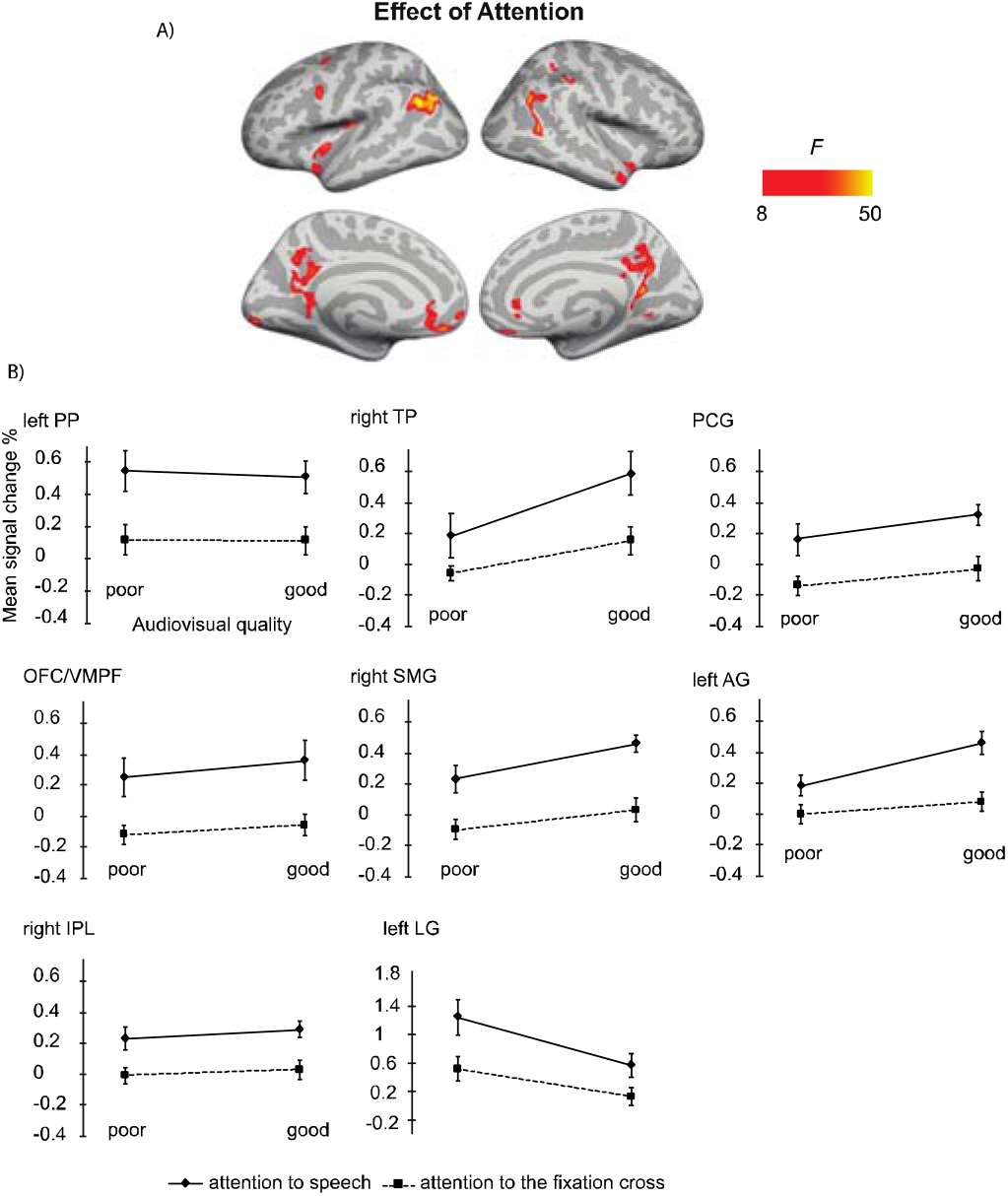
A: Brain areas showing significant main effects of Attention (attention audiovisual speech vs. attention to speech attention to the fixation cross); p(cluster-level FWE) < .05, voxel-wise threshold p <. 001, cluster extent k = 100. B: Mean signal changes (%) compared with rest in the clusters in the showing significantly higher activity during attention to speech than during attention to the fixation cross conditions. No cluster showed an opposite effect. PP, planum polare; TP, temporal pole; PCG, posterior temporal gyrus; OFC/VMPF, orbitofrontal cortex/ventromedial prefrontal cortex; SMG, supramarginal gyrus; AG, angular gyrus; IPL, inferior parietal lobule; LG, lingual gyrus.

## 4. Discussion

We investigated brain areas activated during selective attention to audiovisual dialogues. In particular, we sought attention-related modulations in the auditory cortex and fronto-parietal activity during selective attention to naturalistic dialogues with varying auditory and visual quality. Behaviorally, we observed that increased quality of both auditory and visual information resulted in improved accuracy in answering to the questions related to the content of dialogues. Hence, expectedly, both increased auditory quality (e.g., Davis and Johnsrude, 2003) and increased visual quality (Sumby and Pollack, 1954) facilitated speech comprehension. However, no significant interaction between Auditory Quality and Visual Quality was observed. Thus, our results are not able to give full support to maximal facilitation of speech processing by visual speech at the intermediate signal-to-noise ratio reported, for instance, by McGettigan and colleagues (2012) and Ross and colleagues (2007).

In the fMRI analysis, the main effect of Auditory Quality showed that increasing speech quality was associated with increased activity in the (bilateral) STG/STS, which corroborates previous studies on speech intelligibility (e.g., Scott et al., 2000, Davis and Johnsrude, 2003, Obleser et al., 2007, Okada et al., 2010, McGettigan et al., 2012, Evans et al., 2014, Evans et al., 2016). Enhanced activity in the STG/STS bilaterally is most probably related to enhanced speech comprehension with increasing availability of linguistic information. The STG/STS activity extended to the temporal pole, which might be associated with enhanced semantic processing with increasing speech quality (Patterson et al., 2007). The right STG/STS activity observed here might also be related to prosodic processing during attentive listening (Alho et al., 2006, McGettigan et al., 2013, Kyong et al., 2014). The right temporal pole, in turn, its most anterior part in particular, has been also associated with social cognition (Olson et al., 2013), which may have been triggered by our naturalistic audiovisual dialogues. In addition to the temporal lobe activity, we observed increasing activity in the left angular gyrus and left medial frontal gyrus with increasing speech intelligibility. Enhanced activity in the left angular gyrus may reflect successful speech comprehension, stemming either from increased speech quality or from facilitated semantic processing due to improved speech quality (Humphries et al., 2007, Obleser and Kotz, 2010). The left medial frontal gyrus, in turn, has been attributed to semantic processing as a part of a semantic network (Binder et al., 2009). Hence, an increase in these activations with improving speech quality implies a successful integration of linguistic information onto the existing semantic network and improved comprehension of the spoken input – extending beyond the STG/STS.

The main effect of Visual Quality demonstrated increasing activity in the bilateral occipital cortex and right fusiform gyrus with decreasing visual quality – areas related to object and face recognition (e.g., Weiner and Zilles, 2016). Enhanced activity in these areas might be due to, for instance, noise-modulation of the videos that contained more random motion on the screen than good quality videos. Visual noise has been shown to activate primary regions in the occipital cortex more than coherent motion (e.g., Braddick et al., 2001). It is, however, also possible that viewing masked visual speech required more visual attention than viewing the unmasked videos, causing enhanced activity in the degraded visual conditions. Nevertheless, activity in the middle occipital gyrus was higher for poor visual quality combined with poor auditory quality than for good visual and auditory quality even during attention to the fixation cross (Figure 4). This suggests that increased visual cortex activity for poorer visual quality was at least partly caused by random motion of the masker. Activity enhancements with poor (contra good) audiovisual quality were also observed in the left superior parietal lobule, precuneus and right inferior parietal lobule in both attention conditions, implying contribution of random motion in the masker to these effects as well. Increased visual quality was also associated with enhanced activity in the bilateral STG/STS, corroborating other studies reporting these areas being involved in multisensory integration (e.g., Beauchamp et al., 2004, Beauchamp et al., 2004). We also observed an increase in the left IFG activity with increasing visual quality, an area related to the processing of high-order linguistic information (e.g., Obleser et al., 2007). Enhanced activity in the left IFG has been also observed during integration of speech and gestures (Willems et al., 2009), suggesting its involvement in multimodal integration also in the current study.

We also performed the 2 × 2 ANOVA on brain activity during attention to speech with activity during attention to the fixation cross for stimuli with poor and good audiovisual quality. There, the main effect of Audiovisual Quality in the ANOVA indicated higher bilateral STG/STS (extending to the temporal pole) activity for good auditory and good visual quality than for poor auditory and poor visual quality. The STG/STS effects were also observed during attention to the fixation cross, implying quite automatic speech processing with enhanced audiovisual quality. Furthermore, we found that for poorer audiovisual quality, activity was higher in the left superior parietal lobule, the right inferior parietal, the left precuneus, and bilateral middle occipital gyrus, possibly reflecting automatic processing of facial information.

The main effect of Attention in the 2 × 2 ANOVA indicated enhanced activity during attention to speech in the left planum polare, angular and lingual gyrus, as well as the right temporal pole. We also observed activity in the dorsal part of the right inferior parietal lobule and supramarginal gyrus, as well as in the oribitofrontal/ventromedial frontal gyrus and posterior cingulate bilaterally. One might wonder why attending to the dialogues in relation to attending to the fixation cross was not associated with activity enhancements in the STG/STS as in some previous studies on selective attention to continuous speech (e.g., Alho et al., 2003; Alho et al., 2006). One possible explanation is the ease of the visual control task (i.e. counting the rotations of the fixation cross), eliminating the need to disregard audiovisual speech in the background altogether. This interpretation is also supported by the STG/STS activations observed even during attention to the fixation cross, at least when the audiovisual quality in to-be-ignored speech was good (see Figure 4). Areas in the planum polare have been shown to be associated with task-related manipulations in relation to speech stimuli (Harinen et al., 2013, Wikman et al., 2019).

Auditory attention effects have also been reported outside the STG/STS, for instance, in the middle and superior frontal gyri, precuneus, as well as superior parietal inferior and superior parietal lobule (e.g., Degerman et al., 2006, Salmi et al., 2007). These areas are at least partly involved in the top down control of auditory cortex during selective attention. Interestingly, even though the participants attended to visual stimuli both during attention to speech and attention to the fixation cross, activity was higher in the lingual gyrus (approximately in areas V2/V3 of the visual cortex) during attention to speech. This effect is presumably explained by differences in visual attention between the tasks (see, e.g., Martínez et al., 1999). In other words, while both tasks demanded visual attention, task-related processing of visual speech was presumably more attention-demanding, especially when the faces were masked, as compared to processing of fixation-cross rotations. In line with the previous studies on selective attention to continuous speech (Alho et al., 2003, Alho et al., 2006, Scott et al., 2004), attention to audiovisual dialogues did not significantly engage dorsolateral prefrontal and superior parietal areas. This may be due to high automaticity of selective listening to continuous speech, which might, hence, be quite independent of fronto-parietal attentional control (Alho et al., 2006). However, for the present audiovisual attention to speech, we observed activation in the left inferior parietal lobule, which may be related to attentive auditory processing (e.g., Alain et al., 2010, Rinne et al., 2009).

Furthermore, attention to audiovisual speech elicited enhanced activity in the orbitofrontal/ventromedial prefrontal cortex in comparison with attention to the fixation cross. One possible explanation would be that this activity is related to processing of semantic information (e.g., Binder et al., 2009) in attended speech in contrast to visual information in the fixation cross. Alternatively, this effect may be related to the social aspect of the attended dialogues, since the ventromedial frontal area is associated with social cognition, such as theory of mind and moral judgment (Bzdok et al., 2012), as well as evaluation of other persons’ traits (Araujo et al., 2013). Moreover, enhanced activity in the posterior cingulate and right superior temporal pole observed here during attention to speech may be related to social perception, as both these areas have been involved in social cognition (Bzdok et al., 2012). To our knowledge, no previous study has shown that attending to emotionally neutral dialogues would enhance activity in these three brain regions related to social perception and cognition.

To summarize, our study is the first to present findings on selective attention to natural audiovisual dialogues. Our results demonstrate that increased auditory and visual quality of speech facilitated selective listening to the dialogues, seen in enhanced brain activity in the bilateral STG/STS and the temporal pole. Enhanced activity in the temporal pole might be related to semantic processing particularly in the left hemisphere, whereas in the right hemisphere, it may index processing of social information activated during attention to the dialogues. The fronto-parietal network was associated with enhanced activity during attention to speech, reflecting top-down attentional control. Attention to audiovisual speech also activated the orbitofrontal/ventromedial prefrontal cortex – a region associated with social and semantic cognition. Hence, our findings on selective attention in realistic audiovisual dialogues emphasize not only involvement of brain networks related to audiovisual speech processing and semantic comprehension but, as a novel observation, the social brain network.

## 5. Acknowledgments

This study was supported by the Academy of Finland (grant number 297848). MW was awarded with Erasmus+ scholarship. We thank Artturi Ylinen for help with stimulus preparation and data collection and Miika Leminen for methodological support.

## 6. Author Contributions Statement

AL, KA, MW and VS designed the fMRI experiment; AL, MW and VS prepared the stimuli. MW collected the fMRI data, AL and MW performed the analysis; AL, MW and KA wrote the manuscript in collaboration with PW, WS and MM. MM, VS and PW contributed to the fMRI data analysis.

## 7. Conflict of Interest Statement

The authors declare that the research was conducted in the absence of any commercial or financial relationships that could be construed as a potential conflict of interest.

